# Phylosymbiosis and the hologenome in fungus-gardening ants

**DOI:** 10.64898/2026.08.02.742319

**Authors:** Katherine Beigel, Elizabeth D. Boshers, Blake Bringhurst, Katrin Kellner, Jon N. Seal

## Abstract

Hosts and their microbiomes can be extensively integrated so that they behave across evolutionary scales as a single unit or as a ‘hologenome’. Hosts and their associated microbiomes can potentially exhibit concordance among their respective phylogenetic histories, a phenomenon known as phylosymbiosis. The obligate symbiosis of fungus-gardening ants represents a complex symbiotic system as there are two macroscopic hosts that have associated microbiomes. Here we apply 16S rRNA gene analysis of microbiomes in combination with phylogenetic analysis of nuclear genes and SNPs (single nucleotide polymomorphisms) of hosts to determine the extent to which phylosymbiosis characterizes the coevolution among ant hosts, fungal symbionts and bacterial microbiomes of five species of fungus-gardening ants. The strongest evidence of phylosymbiosis (i.e., congruent topologies between host phylogenetics and microbiome structure) was found between ants and ant-associated microbiomes. While fungal phylogenies and microbiome dendrograms were correlated, these correlations were not topologically congruent. We conclude that phylosymbiosis is present between the ant hosts and their ant-associated microbiome and this would be the strongest evidence of the hologenome concept within the attine symbiosis.

## Introduction

Microbiomes— communities of microbes that inhabit the inside and outside of organisms— have a substantial influence on the physiology and behavior of their hosts (1–3). Microbiomes often display remarkable host specificity (4) with microbial community structure reflecting some function of host biology (5, 6) and this function may be obligate (7). Microbiome-host associations vary in specificity and duration, with some microbiota transiently acquired by a host, and others so entangled with their host that they seem to function as a single interconnected unit termed a “symbiome” or “holobiont” (8–12). Within a holobiont, host and symbiont effectively behave as a single organism (13–15); however the concept of a holobiont is not widely accepted as microbiomes may lack the high partner fidelity required to function as a single selective unit across evolutionary and ecological scales, thus retaining traits more consistent with an ecological community (16–19).

The concept of phylosymbiosis provides a quantitative framework to test the relevance of the holobiont model by exploring the degree of coevolution between microbiomes and host phylogenies (8, 20, 21). Although microbial ecology has made major advances in recent years in describing microbiomes across species and spatial distributions (22–26), phylosymbiosis has been explored comparatively less (27–29). Relatively lacking are phylosymbiotic studies on symbioses that consist of more than one host, which is surprising since symbioses that are characterized by codispersal or coinheritance of symbionts are at least qualitatively consistent with the hologenome concept (30, 31). It would be valuable to understand which parts of a symbiosis are consistent with a hologenome or a community.

Fungus-gardening ants (Myrmecinae, subtribe Attina) have formed a complex macrosymbiosis with specific strains of fungi that they cultivate as an external digestive organ (32). The fungi cultivated by these ants are generally understood to be vertically transmitted through codispersal (33–35). Though vertical transmission is the general rule, horizontal movement of fungal symbionts is not infrequent (36–42). Both fungus-gardening ants and their fungi associate with diverse bacteria and microfungi. These additional microbes are thought to have significant impacts on the overall function of the symbiosis, including moderating pathogen susceptibility, detoxifying harmful compounds, fixing nitrogen, and producing nutrients (43–52). Some attine ant-associated microbiomes display strong host specificity, distinguishable between colonies of a single species and sometimes even among the castes of a single colony (53–56). Moreover, specific taxa within the attine microbiota have cophylogenetic associations, such as Actinomycetota, formerly “Actinobacteria” (45, 57, 58), and Mollicutes(34).

Vertical symbiont transmission likely reinforces phylosymbiotic patterns (59, 60).. For example, comparative studies have found that ant-associated microbiome composition appears to be a function of host species, which is suggestive of vertical transmission via codispersal (53, 56, 61, 62). However, one study concluded that significant components of *Mycocepurus smithii*-associated microbiomes are acquired horizontally from environmental sources (55), as microbiome structure was not correlated with ant or fungal genetics. Relatively few studies have simultaneously evaluated both ant and fungus associated microbiomes, especially in closely related species (55, 61). While one study depicted clear differences in bacterial microbiome structure of two co-occurring ant species (*Trachymyrmex septentrionalis* and *Mycetomoellerius turrifex*), bacterial composition of their fungus gardens appeared to be environmentally determined (Bringhurst et al 2022). Although numerous studies have described the microbiomes of fungus-gardening ants and their fungus gardens, there have been few if any explicit phylosymbiotic tests. Circumstantial evidence of phylosymbiosis seems strongest between the ants and their associated bacteria since the ant host generally seems to exert a strong structuring influence on their microbiota, and weak at best for the fungus garden and its associated bacteria, which lack strong host-structured signals. However, strong host signals (host correlation with microbiome structure) can be potentially confounded by behavior or geography, resulting in the ‘false’ appearance of phylosymbiosis. For example, if hosts differ in behavior that impacts cultivation of bacteria, their microbiomes may also differ, resulting in the appearance of significant host-bacteria phylosymbiosis, even if bacteria were horizontally acquired (Moran and Sloan 2015).

To explore phylosymbiotic structure within the fungus-gardening ant symbiosis, we combined newly acquired and preexisting datasets to evaluate the extent of phylosymbiotic correlations and congruence within the dual host symbioses of five fungus-gardening ant species in the genera *Trachymyrmex* and *Mycetomoellerius*. As these species ants grow related lineages of fungi only known from associations with ‘higher attine’ genera, they should be expected to show extensive phylosymbiosis (32, 63–65). We used a combination of 16S rRNA gene sequencing/analysis, single nucleotide polymorphisms (SNPs) from whole genome sequencing (WGS), and Sanger sequencing of marker genes to 1) characterize the microbiomes associated with ant hosts and fungal symbionts, and 2) measure phylogenetic congruence between ant hosts, fungus symbionts, ant-associated microbiomes, and fungus-associated microbiomes of five North American fungus-gardening ant species. Geographic location was used as an environmental proxy to evaluate the relative impacts of environment and host lineage on microbiome structure.

## 2. Methods

### Sampling methods

Workers and fungus garden material were collected from *Mycetomoellerius turrifex*, *Trachymyrmex septentrionalis*, *Trachymyrmex arizonensis*, *Trachymyrmex pomonae*, and *Trachymyrmex smithi*. See Table S1 for a description and location of data used in this study.

*Trachmyrmex* and *Mycetomoellerius* species are commonly distributed in the southern US and Mexico (66). The five ant species examined in this study were located in two different geographic regions in the southern United States. *M. turrifex* and *T. septentrionalis* were sampled in May/June of 2016 near Bastrop, Texas and Tyler, as described in Bringhurst et al. (2022); these sites are located in the Post Oak Savannah ecoregion, characterized by sandy soils and dominated by oaks and pines. *Trachymyrmex arizonensis* and *T. pomonae* were sampled in July/August of 2018 in the Chiricahua Mountains of the Coronado National Forest near Portal, Arizona (referred to as “Arizona”) as described by Beigel et al. (2021).*Trachymyrmex smithi* was sampled in June of 2022 in the Chihuahua Desert near Horizon City, Texas, USA

Following the excavation of a nest, workers and fungal material were collected from inside nest chambers, from the same chamber whenever possible. Soil samples were collected from near the nest chambers as a negative microbiome control. To minimize contamination, collecting equipment was flame-sterilized with >70% ethanol and resterilized between sample collections. Microbiome samples were preserved in sterile microcentrifuge vials in 100% molecular-grade ethanol and stored at −80 °C.

### Molecular methods

Genomic DNA was extracted from individual workers using the Qiagen QIAamp DNA Micro Kit, following the manufacturer’s protocol with minor alterations. The abdomen was removed from each ant before extraction to reduce potential genomic contamination from gut contents. Two elution buffers were used in the extractions: molecular grade nuclease-free water was used for elution of nuclear gene extracts, while Qiagen Buffer AE was used for elution of the more sensitive SNP extracts.

Three nuclear genes were targeted for PCR amplification, as described in Beigel et al. (2021): elongation factor 1-alpha-F1 (Ef1a-F1), long-wavelength rhodopsin (LW Rh), and wingless (Wg), using primer pairs U52.1(F1-1109F)/L53(F1-1550R), LR143F/LR639ER, and Wg578F/Wg1032R respectively (67–69). PCR products were purified and sequenced at the DNA Sequencing Facility at the University of Texas at Austin in Austin, Texas on an Applied Biosystems 3730 DNA Analyzer and at Eton Bioscience, Inc. in Durham, North Carolina on an ABI 3730xl DNA Sequencer.

16S rRNA gene processing of the ant, fungus, and soil microbiome samples was conducted by MRDNA in Shallowater, Texas, as described by Bringhurst et al. (2022). DNA extraction was performed with the Qiagen DNEasy Powersoil Pro Kit. PCR amplification was performed with the Qiagen HotStarTaq Plus Master Mix Kit, amplifying the V1-V3 hypervariable region of the 16S rRNA gene using primers Gray28F or Gray 27F/Gray519R. PCR products were sequenced with the Illumina MiSeq platform in PEx300 mode. The sequencing provider conducts regular negative controls.

Nuclear gene sequences were subjected to quality checks and trimming with Chromas version 2.6.6 (https://technelysium.com.au/). SNP identification and processing and variant calling was conducted on a subset of *T. arizonensis* and *T. pomonae* ants and fungus as described in Beigel et al. (2021). Whole-genome SNP analysis was chosen because it provides the necessary genetic resolution for distinguishing between the similar fungal symbionts of the study genera in a meaningful way; the routine method of using fungal barcoding genes (e.g., internal transcribed spacer or ITS) results in limited variation and reduced distinguishability (Beigel et al, 2021).

16S rRNA gene sequences were subjected to initial cleanup at sequencing provider MRDNA, where short sequences (<150 bp), sequences with ambiguous base calls, chimeras, sequences with runs exceeding 6 bp, and singleton sequences were removed. Primers were removed and further quality control and ASV (amplicon sequence variant) analyses were conducted in QIIME2-2020.6 (70). In QIIME 2, sequences were demultiplexed with the qiime2 demux plugin, then denoised with the dada2 plugin (71). During denoising, sequences were truncated wherever the average quality score dropped below 20 (around 250 bp), and forward and reverse reads were merged. Taxonomic assignment for each ASV was based on 99% similarity to reference ASV sequences from the SILVA 132_QIIME database (72–74). ASVs associated with mitochondria and chloroplasts were either manually removed or by using the “qiime taxa filter-table” and “qiime taxa filter-seqs” commands. Sequences from all five species were combined and rarefied to 1280 reads to create the primary combined dataset. An additional data subset was created by combining sequences from only *T. arizonensis* and *T. pomonae* and rarefying to 6900 reads as this two species dataset was of higher quality. The primary ASV table was created from the tabulated taxonomic bar plot of the combined, rarefied dataset of all five species and was analyzed for phylosymbiosis with the ant nuclear markers, while the secondary ASV table was created from the tabulated taxonomic bar plot of the combined, rarefied data subset of *T. arizonensis* and *T. pomonae*, and was analyzed for phylosymbiosis with the ant and fungus SNP markers. The increased read rarefaction threshold for the secondary ASV table was used since both *arizonensis* and *pomonae* ant and fungi samples had higher number of reads than the other species in the nuclear marker analysis, with a read rarefaction threshold of 6900 providing a better community resolution for these two species.

### Statistical analysis

**Microbiome characterization.** Variation in microbial community structures (ASV tables) of each species and each sample type (ants and fungus) was explored using Analysis of Similarities (ANOSIMs) in the R package vegan 2.6-2. Non–metric multidimensional scaling (NMDS) plots were used to visualize these microbial community structures (Oksanen et al., 2018), as microbiomes must be distinct between groups of interest for any downstream analysis to be meaningful (Lim et al., 2020). To determine significant differences in community diversity between groups, mean Shannon’s entropy values were compared with between-sample type and within-sample type with Kruskal-Wallis rank sum tests in R. To determine the factors driving any significance, non-parametric Dunn’s tests (R, FSA 0.9.4) were used as post-hoc pairwise tests where applicable with Bonferroni corrections to adjust p-values (75). The total read counts for each taxon in a group were calculated as a percentage of total read counts of all taxa in that group to summarize the most abundant microbiota across different groupings.

**Evaluation of environmental components of microbiome structure**. To identify the most influential bacterial taxa structuring the microbiomes of each region, indicator species analyses (ISA)_were carried out on ASV relative abundances between east and west for both ant-associated and fungus-associated microbiomes (76). Indicator species analyses were carried out with the indicspecies R package version 1.8.0 (with 9,999 permutations).

To evaluate the overall structure of environment on individual microbial taxa, geographic location was used as a proxy for environment in redundancy analyses (RDA). The dataset was separated into species sampled from two broad geographic regions: ‘East’ (*M. turrifex* and *T. septentrionalis*) and ‘West’ (*T. arizonensis*, *T. pomonae*, and *T. smithi*). Redundancy analyses using the vegan 2.6-2 R package comparing the impact of host ant species and broad geographic region on each influential microbial class (reduced model with 10,000 permutations). Additionally, we explored the impact of geographic location on microbiome structure using partial Mantel tests. These tests quantify the correlation between two factors while accounting for the effects of a third factor. Therefore, correlations between hosts and microbiomes can be quantified while controlling for the effects of geography and correlations between geography and microbiomes can be quantified while controlling for the effects of the hosts. Geographic dissimilarity matrices were created for each region by calculating pairwise geographic distances based on GPS coordinates of each sample with distGeo in R (77); geosphere, v 1.5-18) (78), which uses ellipsoid calculations to ensure greater accuracy for distance measurements (25). If GPS coordinates of an individual colony’s collection site were not available, GPS coordinates of the colony’s collection locale were used instead. Microbiome dissimilarity matrices were composed of weighted UniFrac values calculated in QIIME 2 from 16S rRNA gene data, and host dissimilarity matrices were composed of host phylogeny cophenetic distances calculated with the stats package (v4.2.1) in R. All R analyses were performed using R (79). Separate sets of correlations were conducted for each geographic region. For each region, each microbiome dissimilarity matrix was correlated separately with the host dissimilarity matrix and the geographic distance matrix, while controlling for the effects of the other matrix.

**Phylosymbiosis analyses.** Two tests for phylosymbiosis (genetic distance matrix comparisons and topological congruence tests) were conducted; either test can detect phylosymbiosis but using both metrics reduces the chance of making a Type I error (21). Two sets of these tests were conducted at different resolutions of sequencing data. One set of tests used nuclear gene data from the ant hosts of all five species, paired with the 16s rRNA datasets from all five species; and the other set of tests was conducted using SNP data from both ant hosts and fungus hosts from a subset of colonies sampled in Arizona (12 *T. arizonensis* colonies and 6 *T. pomonae* colonies), paired with the 16s rRNA data subsets of *T. arizonensis* and *T. pomonae*. The SNP dataset depicted several significant clades within ants and fungi that the five species nuclear marker dataset was unable to detect, providing a high-resolution tool for exploring intraspecific patterns of phylosymbiosis, although less taxonomically broad than the five species nuclear marker dataset.

Genetic distance matrix correlations compare the genetic distance matrix of a microbiome with that of the host, detecting phylosymbiosis if host and microbiome matrices correlate (21). Microbiome genetic distance matrices for ant-associated and fungus-associated microbiomes were created by calculating weighted UniFrac values from 16s rRNA gene data in QIIME 2. The ant host genetic distance matrix for all five species was created by calculating the cophenetic distances (R stats, v4.2.1) of the nuclear gene multi-gene phylogenetic tree in R, resulting in a matrix of cophenetic distances between all taxa. The ant host and fungal host genetic distance matrices for the Arizona subanalysis were created by calculating the Euclidean distances between each SNP sample based on the genotype at each SNP locus in R (adegenet, v2.1.8; stats, 4.2.1) (80, 81).

Microbiome distance matrices were correlated with host matrices using Procrustes superimposition in R (vegan v2.7-2), employing PROTEST with 999 permutations for significance testing. Procrustes superimposition calculates the number of modifications necessary to maximize symmetry between two genetic distance matrices, and PROTEST measures the significance of the correlation between the two modified matrices, with possible PROTEST results ranging from 0 (no correlation) to 1 (complete correlation).

Topological congruence tests compare a microbiome dendrogram and host phylogenetic tree, detecting phylosymbiosis if the phylogenies are congruent (Lim et al, 2020). Microbiome dendrograms for ant-associated and fungus-associated microbiomes were calculated using 16S rRNA gene data with UPGMA (phangorn v1.12.2) in R (82, 83).

In the five ant species phylogenetic analysis, edited sequences of the three nuclear genes (Ef1a-F1, LW Rh, and Wg) were trimmed for quality control and *Sericomyrmex parvulus* was used as the outgroup (voucher USNM JSC051026-01-LS12; NCBI GenBank accessions: MK600360.1 (Ef1a-F1), MK600174.1 (LW Rh), MK600281.1 (Wg)). Sequences were aligned for each gene using ClustalW in MEGA11 (84). Aligned sequences for each gene were concatenated using AMAS (v1.00) (85). The most suitable substitution model for each gene was identified using ModelTest-NG (v0.2.0) (86, 87). Maximum likelihood (ML) phylogenetic inference was performed using RAxML-NG (v1.2.2) (88). ML tree searches were conducted from 1,000 starting trees (500 parsimony-based trees and 500 random trees) using DNA partitioning to apply the substitution model identified by ModelTest to each gene in the multiple sequence alignment. Node support was assessed with 1,000 nonparametric bootstrap replicates using the best-fit model inferred during the ML tree search.

Ant host and fungus host phylogenies from Beigel et al. (2021) were used for the Arizona species subanalysis. Briefly, phylogenies were calculated using whole genome sequence SNPs using a published genome of *T. septentrionalis* and a published draft genome of a fungus extracted from a *T. arizonensis* colony (36, 89). Low coverage genomes were obtained from prepared genomic DNA extracts and sequenced on an Illumina NovaSeq 6000 sequencer that generated 150 bp paired end reads at SNPsaurus (Eugene, Oregon). Initial bioinformatics (demultiplexing, alignment variant calling, and SNP filtering) was conducted by SNPsaurus. Demultiplexing, trimming and mapping to the reference genome were accomplished using BBTools (*bbduk*, *bbmap*) (90). Variants were called using *callvariants.sh* (within BBMap) of aligned reads with a minimum mapping quality of 15. VCFTools was used to filtered called vari-ants (minimum depth of 8 and minimum allele frequency of 5%). (91). After alignment with the reference genome and the removal of invariant sites, the ant and fungal SNP alignments contained approximately 1.5 million bp and 90,000 bp, respectively. In RAxML (v8.2.11), 1,000 independent maximum likelihood tree searches were performed. RAxML was run using the recommended GTR+GAMMA model for SNP data (ASC_GTRGAMMA) and the Felsenstein method for ascertainment bias correction (92–94). Branch support for the best-scored tree was evaluated using bootstrap analysis with 1,000 replicates. (92–94)

Phylosymbiosis requires that the topology of the microbiome dendrogram is congruent with the topology of the host phylogenetic tree (21). Congruencies between the microbiome dendrograms and the host phylogenetic trees were checked for phylosymbiotic significance using a Python script from Brooks et al. (2016) to calculate normalized Robinson-Foulds and normalized matching cluster values with a minimum of 10,000 random trees. Both Robinson-Foulds and matching cluster metrics evaluate congruence between two topographies, with possible results ranging from 0 (complete congruence) to 1 (incomplete congruence) (9).

Robinson-Foulds finds the smallest number of operations necessary to perfectly match one topology to the other, while matching cluster takes subtree-level congruence into account, resulting in a more refined evaluation of small topological changes; accordingly, the Matching Cluster metric is a better evaluator of fine sub-tree level differences (21).

## 3. Results

### Microbiome characterization

**Ant-associated microbiomes**. Ant-associated microbiomes were unique to each species and were generally different from the fungus-associated microbiomes (Figure 1; Table 1,2; Figure S1). The single exception was that *T. arizonensis* ant microbiomes were not different from the bacterial microbiomes of their fungus gardens when included in the multispecies dataset; however, *T. arizonensis* ant microbiomes were significantly different from *T. pomonae* microbiomes in the analysis of Arizona-only dataset (see Methods; Table 1). Overall microbial community alpha diversity was significantly different among the ant-associated microbiomes (Kruskall-Wallis test using mean Shannon’s entropy values: df =4, χ2=11.191, p-value=0.0245; Figure S2, Table S2). The bacterial community of each ant species appeared to be composed of a different set of bacterial taxa rather than subsets (e.g., ASVs) of the same bacterial taxa (Figure 1; Figure S2, Table S2).

**Table 1.**
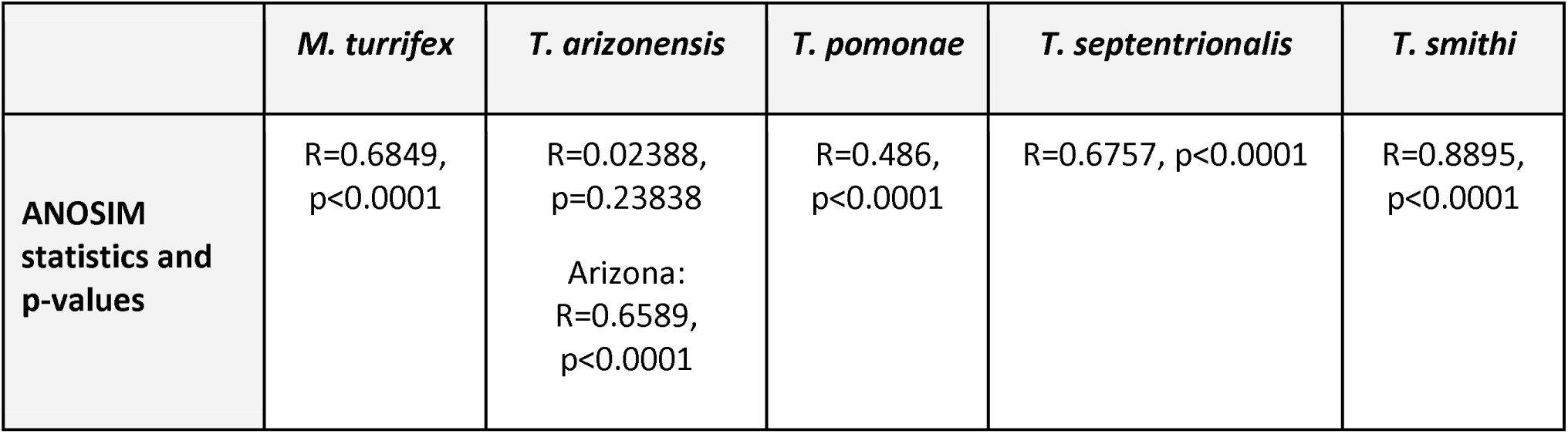
ANOSIM results between the ant-associated microbiome and fungus-associated microbiome for each species. Bacterial microbiomes of each ant species are different from the microbiomes associated with the fungus grown by each species. Bacterial microbiomes of ants and fungi of T. arizonensis were not significantly different in the five species dataset, but were different in the Arizona-only dataset (T. arizonensis and T. pomonae).

**Table 2.** ANOSIM results comparing ant-associated microbiomes of each species. All ant-associated microbiomes were significantly different from each other.

| **ANTS** | ***M. turrifex*** | ***T. arizonensis*** | ***T. pomonae*** | ***T. septentrionalis*** |
| --- | --- | --- | --- | --- |
| ***T. arizonensis*** | R=1, p<0.0001 |  |  |  |
| ***T. pomonae*** | R=0.8448,<br>p<0.0001 | R=0.8489,<br>p<0.0001 |  |  |
| ***T. septentrionalis*** | R=0.8189,<br>p<0.0001 | R=1, p<0.0001 | R=0.859,<br>p<0.0001 |  |
| ***T. smithi*** | R=0.9613,<br>p=0.0011 | R=1, p=0.0004 | R=0.8593,<br>p=0.0007 | R=1, p=0.0003 |

The ant-associated microbiome of *T. arizonensis* was dominated in abundance by two main taxa: *Pseudomonas* (56.58% of all *T. arizonensis* ant-associated microbiome taxa), and *Planococcus* (25.61%); the ISA identified *Pseudomonas*, unidentified Firmicutes, and unidentified Bacteria as taxa significantly associated with *T. arizonensis* (Figures 1 and 2).

The ant-associated microbiome for *T. pomonae* was dominated in abundance by strains of unidentified Intrasporangiaceae (16.52% of all *T. pomonae* ant-associated microbiome taxa), *Planococcus* (15.77%), *Nocardioides* (14.40%), and *Spiroplasma* (13.28%). The ISA identified *Nocardioides, Marinilutecoccus*, unidentified Actinomycetota and Solirubrobacteraceae, *Spiroplasma*, and *Nocardia* as significantly associated with *T. pomonae* (Figures 1 and 2).

The ant-associated microbiome of *T. smithi* was dominated by *Aeromicrobium* (55.06% of all *T. smithi* ant-associated microbiome taxa) and *Wolbachia* (16.91%), with *Aeromicrobium*, *Brevibacterium*, unidentified Pseudonocardiaceae, *Wolbachia*, and *Pseudonocardia* found as significantly associated with *T. smithi* (Figures 1 and 2).

The ant-associated microbiome of *T. septentrionalis* was dominated in abundance by strains of *Solirubrobacter* (34.44% of all *T. septentrionalis* ant-associated microbiome taxa) and *Luteimonas* (14.51%), with two strains of *Solirubrobacter, Aeromicrobium, Microlunatus, Luteimonas*, and *Lautropia* found as significantly associated with *T. septentrionalis* (Figures 1 and 2).

The ant-associated microbiome of *M. turrifex* was dominated in abundance by *Amycolatopsis* (19.87%) and unidentified Burkholderiaceae (14.75%); the ISA identified *Amycolatopsis, Luteipulveratus mongoliensis, Cloacibacterium, Microbacterium, Stenotrophomonas, Pseudonocardia*, unidentified Microbacteriaceae, *Pedobacter*, Methylophilaceae, *Staphylococcus hominis novobiosepticus*, and unidentified Sphingomonadaceae as taxa significantly associated with *M. turrifex* (Figures 1 and 2).

The ISA also identified taxa significantly associated for several species groups: *T. arizonensis* and *T. pomonae* (*Planococcus*), *T. septentrionalis* and *M. turrifex* (unidentified Burkholderiaceae), *T. pomonae* and *T. septentrionalis* (unidentified Intrasporangiaceae), and a triad group of *T. pomonae*, *T. septentrionalis*, and *T. smithi* (*Naumannella*) (Figures 1 and 2).

**Fungus-associated microbiomes.** The fungus-associated microbiomes of each species were generally unique: microbial community structure of the fungus-associated microbiome of each species was significantly different from the fungus-associated microbiomes of the other species (ANOSIMs, p values =<0.005, Table 3), with a single exception: the *M. turrifex* fungus-associated microbiome was not significantly different from the *T. septentrionalis* fungus-associated microbiome (Bringhurst et al. (2022)) (Table 3). Overall microbial community alpha diversity did not vary significantly between fungus-associated microbiomes of different ant species (Kruskall-Wallis results using mean Shannon’s entropy values: df=4, χ2=6.091, p-value=0.1925; Figure S3).

**Table 3.** ANOSIM results between fungus-associated microbiomes of each species. Significantly different comparisons are indicated with a *.

| **FUNGUS** | ***M. turrifex*** | ***T. arizonensis*** | ***T. pomonae*** | ***T. septentrionalis*** |
| --- | --- | --- | --- | --- |
| ***T. arizonensis*** | R=0.7152,<br>p<0.0001* |  |  |  |
| ***T. pomonae*** | R=0.466,<br>p<0.0001* | R=0.7797,<br>p<0.0001* |  |  |
| ***T. septentrionalis*** | R=0.01041,<br>p=0.34 | R=0.6893,<br>p<0.0001* | R=0.5787,<br>p<0.0001* |  |
| ***T. smithi*** | R=0.4613,<br>p=0.0017* | R=0.9601,<br>p=0.0005* | R=0.6426,<br>p=0.0015* | R=0.3812,<br>p=0.008* |

The fungus-associated microbiome of *M. turrifex* was dominated by *Spiroplasma* (21.27% of all *M. turrifex* fungus-associated microbiome taxa) and *Mesoplasma* (13.37%) (*Edwardiiplasma* (95); with unidentified Kryptoniales, unidentified Pyrinomonadaceae, *Ralstonia*, *Thermicanus*, Omnitrophicaeota, and Ochrobactrum found as significantly associated with *M. turrifex* (Figures 1 and 2).

The fungus-associated microbiome of *T. arizonensis* was dominated by the same two species that dominated the species’ ant-associated microbiome: *Pseudomonas* (65.48% of all *T. arizonensis* fungus-associated microbiome taxa) and *Planococcus* (16.22%); *Pseudomonas*, *Planococcus* sp. HL01, *Pseudomonas* sp. BSL2A, *Kurthia* sp. MBG49, and unidentified Gammaproteobacteria were found as significantly associated with *T. arizonensis* (Figures 1 and 2).

The fungus-associated microbiome of *T. pomonae* was dominated by *Spiroplasma* (48.30% of all *T. pomonae* fungus-associated microbiome taxa) and *Pseudomonas* (10.05%); *Spiroplasma, Crossiella*, unidentified Gemmataceae, *Acidothermus*, unidentified Xanthobacteraceae, unidentified Acidobacteriales, *Pseudonocardia* sp. CC980219.11, Ruminococcaceae NK4A214.group, *Gemmata*, and *Chloroflexia* were found significantly associated with *T. pomonae* (Figures 1 and 2).

The fungus-associated microbiome of *T. septentrionalis* was dominated by *Mesoplasma* (41.88% of all *T. septentrionalis* fungus-associated microbiome taxa) and *Spiroplasma* (9.60%); *Bradyrhizobium* and unidentified Tepidisphaerales were found significantly associated with *T. septentrionalis* (Figures 1 and 2). The fungus-associated microbiome of *T. smithi* was dominated by *Mesoplasma* (19.12% of all *T. smithi* fungus-associated microbiome taxa) and unidentified Bacteria (12.17%); unidentified Pseudonocardiaceae, *Blastocatella*, *Methylobacterium*, *Curtobacterium* sp. LJ21, *Nocardioides*, unidentified Caulobacteraceae, unidentified Actinomycetota, *Massilia*, *Nocardioides*, *Enhydrobacter*, *Pelomonas*, Elusimicrobia, and unidentified Sphingobacteriales were found significantly associated with *T. smithi* (Figure 1 and 2).

The ISA also identified several taxa significantly associated with species groupings: *T. arizonensis* and *T. smithi* (unidentified Bacteria), and *T. septentrionalis* and *M. turrifex* (*Solirubrobacter*, unidentified Burkholderiaceae, *Luteimonas*, and *Acidobacteria*) (Figure 2).

**Impact of location on microbiomes.** The partial Mantel tests suggest that ant-associated microbiomes were most influenced by host ant species, while the fungus-associated microbiomes are influenced by a combination of host ant species and environment (in the western fungus-associated microbiomes) or not significantly by either factor (in the eastern fungus-associated microbiomes). The partial Mantel tests found that, in the eastern species (*T. septentrionalis* and *M. turrifex*) host ant species provided the greatest contribution to ant-associated microbiome structure (partial Mantel test between ant microbiome and ant phylogeny when controlling for geography, p = 0.0180, partial Mantel test between ant microbiome and geography when controlling for ant host phylogeny, p = 0.635). Fungus-associated bacterial microbiome structure was not significantly impacted by either host ant species or geographic location (partial Mantel test between fungus microbiome and ant host phylogeny when controlling for geography, p = 0.3280; partial Mantel test between fungus microbiome and geography when controlling for ant host phylogeny, p = 0.3320). Among the western species, (*Trachymyrmex arizonensis*, *T. pomonae*, and *T. smithi*), host ant species also provided the greatest contribution to ant-associated microbiome structure (partial Mantel test between ant microbiome and ant phylogeny when controlling for geography, p = 0.001; partial Mantel test between ant microbiome and geography when controlling for host ant phylogeny, p = 0.207), while fungus-associated bacterial microbiome structure was significantly impacted by both host ant species and geographic location (partial Mantel test between fungus microbiome and ant phylogeny when controlling for geography, p = 0.001; partial Mantel test between fungus microbiome and geography when controlling for host ant phylogeny, p = 0.001).

The redundancy analysis (RDA) indicates variable drivers are influencing microbial taxa within microbiomes: some taxa are influenced by host ant species, some by environment, and some by a combination of both factors. The redundancy analyses found that within the ant-associated microbiome, four taxa (Bacteroidia, Verrucomicrobiae, Acidobacteriia, and Mollicutes) were structured by region, one (Alphaproteobacteria) by host ant species, and four (Actinomycetota, Thermoleaphilia, Bacilli, and Gammaproteobacteria) by an interaction of region and host ant species (Table 4). Within the fungus-associated microbiome, three taxa (Thermoleophilia, Bacteroidia, and Verrucomicrobiae) were structured by region, and four taxa (Actinomycetota, Gammaproteobacteria, Mollicutes, and Bacilli) were structured by a combination of region and host ant species (Table 4).

**Table 4.** Results of redundancy analysis (RDA) conducted on the major classes of microbiota identified by the ASV-level ISA comparing the impact of host ant species and geographic region for the ant-associated microbiome and fungus-associated microbiome. All results had p-values < 0.05.

|  | Region | Host ant species | Region and host ant species |
| --- | --- | --- | --- |
| **Ant-associated microbiomes** | Bacteroidia (F=2.8700) | Alphaproteobacteria (F=5.71) | Actinomycetota (region: F=9.4928; species: F=17.0244) |
|  | Verrucomicrobiae (F=1.2768) |  | Thermoleophilia (region: F=93.166; species: F=49.692) |
|  | Acidobacteriia (F=1.7908) |  | Bacilli (region: F=22.5846; species: F=3.9544) |
|  | Mollicutes (F=2.4932) |  | Gammaproteobacteria (region: F=27.899; species: F=19.002) |
| **Fungus-associated microbiomes** | Thermoleophilia (F=8.0076) | None | Actinomycetota (region: F=2.0568; species: F=5.8280) |
|  | Bacteroidia (F=5.6496) |  | Gammaproteobacteria (region: F=15.300; species: F=13.795) |
|  | Verrucomicrobiae (F=2.2419) |  | Mollicutes (region: F=5.7257; species: F= 3.4739) |
|  |  |  | Bacilli (region: F=10.6360; species: F=8.1176) |

### Phylosymbiosis analyses

#### Ants

Correlations between the genetic distance matrices of ant microbiomes and ant/fungus hosts were significant within the combined dataset of all five species as well as within the higher-resolution *T. arizonensis* and *T. pomonae* subanalysis. In all five species, ant-associated microbiomes significantly correlated with ant host nuclear variation (PROTEST = 0.4016, p = 0.001; Figure 2A). In the *T. arizonensis* and *T. pomonae* subanalysis, ant-associated microbiomes significantly correlated with both ant host SNPs (PROTEST = 0.713, p = 0.001; Figure 2C) and fungus host SNPs (PROTEST = 0.8048, p = 0.001; Figure 2E).

Topological congruence comparisons between the ant-associated microbiomes and ant hosts were significant in the combined dataset of all five species (nMC = 0.4725 (normalized Matching Cluster), p = 0; nRF = 0.8800 (normalized Robinson-Foulds), p = 0.001; Figure 2A) as well as for the *T. arizonensis* and *T. pomonae* subanalysis (nMC = 0.40, p = 0.00025; nRF = 0.81, p = 0.00154; Figure 2C). Topological congruence comparisons between the ant-associated microbiomes and fungus hosts were not significant (nMC = 0.58, p = 0.41022; nRF = 1.0, p = 1.0) (Figure 2E). These significant correlations and congruencies support phylosymbiosis between the ant-associated bacterial communities and the ant hosts. Phylosymbiosis between fungal hosts and microbiomes was explored only within the *T. arizonensis* and *T. pomonae* subanalysis since only these two species had corresponding fungal intraspecific phylogenies available (36); while genetic distance matrix comparisons were significant, topologies were not congruent, therefore, it is unlikely that phylosymbiosis is occurring between ant-associated bacterial communities and fungus hosts.

#### Fungus

Correlations between the genetic distance matrices of fungus-associated microbiomes and ant/fungus hosts were significant within the combined dataset of all five species as well as within the higher-resolution *T. arizonensis* and *T. pomonae* subanalysis. In the combined five species analysis, the genetic distance matrices of fungus-associated microbiomes correlated significantly with ant hosts (PROTEST = 0.4374 p = 0.02) (Figure 2B). In the *T. arizonensis* and *T. pomonae* subanalysis, fungus-associated microbiomes correlated significantly both with ant hosts (PROTEST = 0.6502, p = 0.001; Figure 2D) and fungus hosts PROTEST = 0.6585, p = 0.017) (Figure 2F).

Topological congruence comparisons between the fungus-associated microbiome and ant hosts was not unambiguously supported in either the combined dataset of all five species (nMC = 0.5963, p = 0.0038; nRF = 0.9600, p = 0.1613; Figure 2B) or in the *T. arizonensis* and *T. pomonae* subanalysis (nMC = 0.51, p = 0.0536; nRF = 0.93, p = 0.1747) (Figure 2D). Though the Matching Cluster analysis was statistically significant in the five-species dataset, the values were >0.5 which suggests incomplete congruence. No significant topological congruence was found between the fungus-associated microbiome and the fungus hosts in the *T. arizonensis* and *T. pomonae subanalysis* (nMC = 0.69, p = 0.9722; nRF = 1.0, p = 1.0) (Figure 2F). Despite the significant genetic distance matrix correlations, the lack of consistently significant topological congruencies suggests that phylosymbiosis is unlikely to exist between the fungus-associated bacterial communities and either the ant or fungus hosts.

## 4. Discussion

The strongest evidence of phylosymbiosis within the fungus-gardening ant symbiosis exists between the ant hosts and ant-associated bacteria. Although this study reports significant genetic and microbiome matrix correlations with the bacterial microbiome of fungus gardens, topology tests only supported phylogenetic congruence consistently between ant hosts and ant-associated microbiomes. Accordingly, the hologenome concept is a potential descriptor only of the ants and at least some of their associated bacteria, as only that host-microbiome pairing appears to have a concordant evolutionary history.

Phylosymbiosis between the fungus garden and its associated bacteria appears much more limited. Even though bacterial microbiomes of fungus gardens were distinct among host ant species, this did not appear to be driven by a co-evolutionary process. Although fungus-gardening ants typically show a mixture of vertical and horizontal transmission of fungal symbionts that suggests ‘diffuse’ coevolution (32, 42) *T. arizonensis* and *T. pomonae* each appear to associate with distinct fungal lineages, which is suggestive of a ‘one-to-one’ coevolutionary relationship (36). Nevertheless, neither the ants nor the fungi grown by these two species appear to be co-diverging with the bacterial communities found in fungus gardens (Figure 2C-E). The fungus garden may be more of a co-host as it is likely heavily influenced by the ants depositing foraged substrate items among the fungal mycelia (96), depositing Actinomycetota, Mollicutes, and other bacterial taxa via fecal droplets (62) and acquiring other bacteria from the environment. Frequent switching of fungal symbionts (42) could also be a mechanism to break up any long term phylosymbiotic relationship with garden bacteria.

As this study demonstrates phylosymbiosis between ants and their bacteria, the most productive next step will be to explore the interactions within the ant-associated bacterial community. Symbiotic functions have been identified for some bacterial groups, especially with regard to those with defensive functions such as the Actinomycetota (*Pseudonocardia*, *Amycolatopsis*, etc.) (57). Somewhat unexpected, the Actinomycetota exhibit a host-environment interaction so even though some Actinomycetota share cophylogenetic history with ants, there is some evidence for environmental uptake. Mollicute bacteria are thought to have crucial nutritional functions by being involved in carbon and nitrogen cycling (97). Some studies have reported colony and species level specificity in Mollicutes, which is consistent with vertical transmission (98, 99); however, the results in this study imply that Mollicutes could be environmentally acquired by the ants (Table 4). Similarly, Mollicutes may be more common in some lineages of fungus gardens than others (100). In any case, except for a single *Spiroplasma* ASV associated with *T. pomonae*, Mollicutes do not appear to be a major contributor toward the significant ant host effects (Figure 1). Although the lack of phylosymbiotic signal with Mollicutes might be an artifact of the relatively short reads used in this study; invariable read length may also indicate relatively recent acquisition and expansion of bacteria throughout host populations. While a phylogenetic analysis of Mollicute bacteria was able to place *Mesoplasma* associated with fungus-gardening ants in a separate clade from *Mesoplasma* associated with distantly related army ants, there was low support within fungus-gardening ants (34). Several other taxa such as those in the Gammaproteobacteria and Alphaproteobacteria appear to have strong host effects (significant Mantel tests and RDA analyses), which could reflect a long evolutionary history. Future work could address whether any of these are simply commensals or provide some crucial role to the symbiosis by incorporating longer read lengths, transcriptome analysis or designs that would target some of these bacteria and test explicit evolutionary questions.

A surprising finding in this comparative study was the restricted distribution of *Solirubrobacter*, which is a characteristic bacterium of *T. septentrionalis* (54, 61). Except for an ASV of Solirubrobacteriaceae found in *T. pomonae* ants, *Solirubrobacter* is restricted to *T. septentrionalis* and thus probably is acquired through environmental uptake. Curiously, *T. septentrionalis* ants contain two distinct ASVs of *Solirubrobacter*, whether these two variants are biologically distinct is not known. Though its function in ants is not understood, it appears to be an important bacterium associating with plant roots in arid environments (54, 61, 101).

One unexpected finding in this study were the correlations between host ant species and bacterial microbiome of fungus gardens. Bringhurst et al. (2022) found that the environment rather than host species explained the most differences in fungus-associated bacterial microbiome structure of *M. turrifex* and *T. septentrionalis*. However, our partial Mantel tests, which included the data of Bringhurst et al. (2022), found that both environment and host species exerted a significant structuring influence on fungus-associated microbiomes. Ant host species was a significant influence in the structure of fungus-garden bacterial microbiomes in all species except *M. turrifex* and *T. septentrionalis* – why these two species are the exception is unclear, although it may be related to differences in sampling. Bringhurst et al. (2022) explored the bacterial microbiomes of colonies collected over a large range (>300km between the most distant sites) whereas colonies of the other species included in this study were obtained from fewer sites within a much narrower geographical range. The greater environmental differences within the *M. turrifex* and *T. septentrionalis* colonies could obscure host-driven patterns because microbiomes containing bacterial taxa acquired from a greater diversity of surrounding soils and ant-collected substrate items may appear less similar.

This study has demonstrated that the presence and strength of phylosymbiosis is not ubiquitous among ant hosts and their microbiota. Weak phylosymbiosis has been reported between birds and their associated bacteria (102), as well as between corals and their photosynthetic dinoflagellates (103) and considerable variation among corals and other bacteria (104). Stronger phylosymbiosis has been reported between corals and their symbiotic bacteria (28) as well as in a variety of other insects and mammals (9, 19, 20, 105). The similarity of microbiomes in many geographically separate insect-fungal symbioses suggest that convergent evolution also represents another possible mechanism for phylosymbiosis (12, 16, 18, 20, 106, 107). Nevertheless, identifying phylosymbiotic partners represents an initial step in understanding holobiont stability as these reflect some importance in their long-term co-evolutionary history.

## Data Availability

The 16S rRNA gene sequences from *T. smithi* are available under NCBI BioProject PRJNA1090512; the 16S rRNA gene sequences from *T. arizonensis* and *T. pomonae* are available under NCBI BioProject PRJNA713779; and the 16S rRNA gene sequences from *T. septentrionalis* and *M. turrifex* are available under BioProject PRJNA789907 (Table S1). The Sanger sequences are available under GenBank Accession Numbers PX132565-PX132593, PX270874-PX270887, MW358558–MW358601, MW694488–MW694712, and MZ050065–MZ050068. WGS data for *T. arizonensis* and *T. pomonae* is available under BioProject PRJNA713779 and the derived SNP data is available on Dryad (https://doi.org/10.5061/dryad.d2547d83m). The ASV tables, Qiime2 pipeline, multi-gene alignments of Sanger sequences, and the Sanger-based phylogenetic tree can be found on Dryad Digital Repository (https://datadryad.org/share/LINK_NOT_FOR_PUBLICATION/i-ErCeBIy3QLdyBoQgk_IOSZpUhmV75Lav9BS9ykZSo). Code for phylogenetic and phylosymbiosis analyses is available at https://github.com/kbeigel/phylosymbio-kit, release tag v1.0.0.

## Acknowledgements

Ethan Van Arnam, Paul Lenhart, and Ulrich Mueller provided valuable advice for field collection of *T. smithi*, and Christine Bays helped with collection of *T. smithi* colonies. Matthew Richards-Perhatch V assisted with organizing bacterial abundances datasheets in Excel, provided technical and creative support for R analyses, and assisted with development of microbiome figures. Mingna Zhuang at the University of Texas at El Paso as well as Valdime Walker and Schi-Lee Smith at the University of Texas at Tyler provided logistical support for *T. smithi* collection. This project was supported by a National Science Foundation CAREER award to JNS (IOS-1552822) and DEB-**2230334 to JNS and KK.** EB was supported by REPS (Research for Post Baccalaureate Scholar) supplement IOS-2136147.

## Author Contributions

JNS and KK conceived the study and obtained financial support. EB and JNS secured permits and EB and BB conducted the fieldwork novel to this study. EB and KB conducted the molecular work and analyzed the nuclear gene sequences. EB and BB analyzed the 16S rRNA gene sequences. KB and EB conducted the statistical analyses. KK, JNS, BB, and KB provided preexisting data. JNS led manuscript writing, assisted by BB, KB and KK. All authors edited and approved the final manuscript.

---

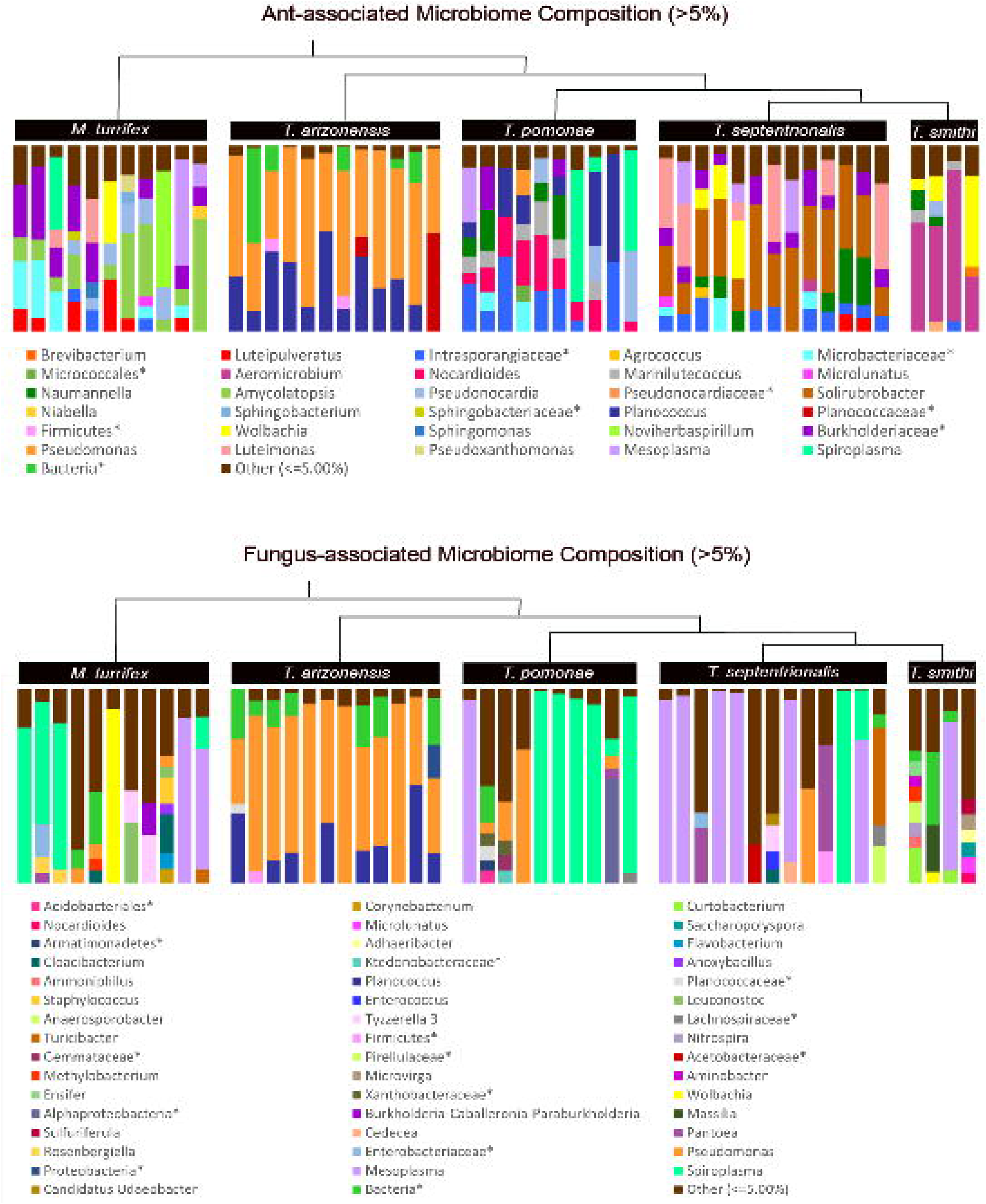

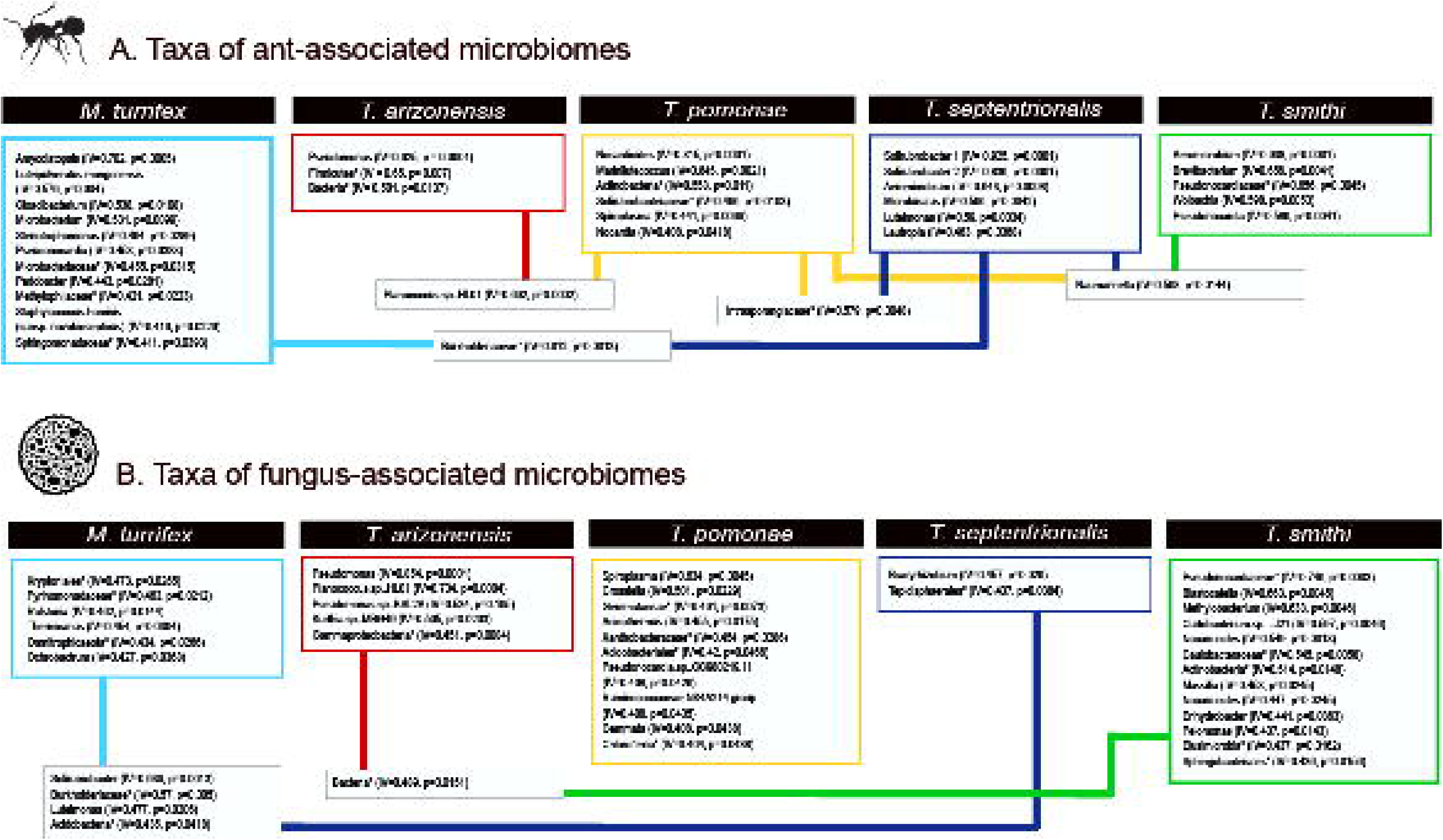

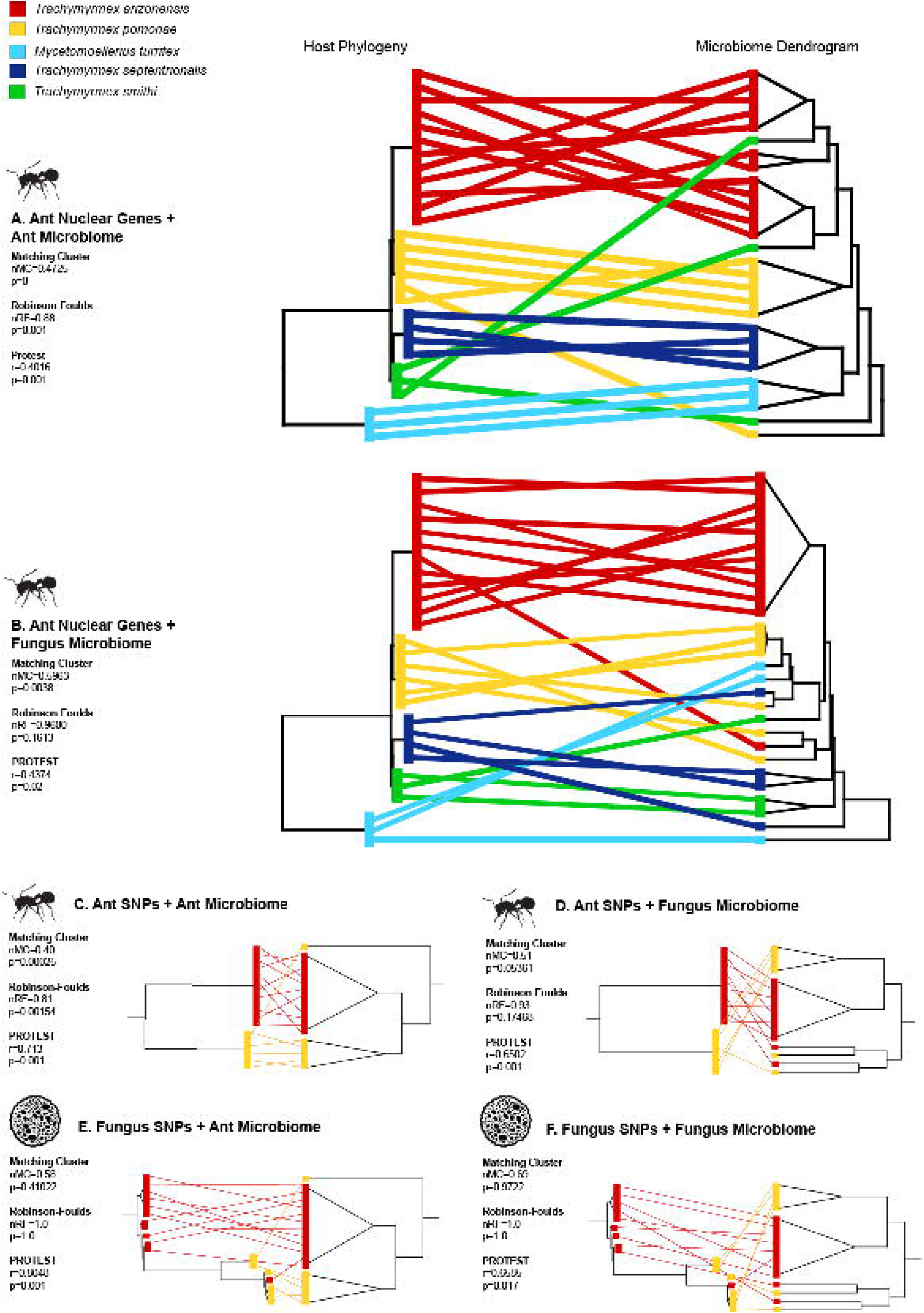

## Notes

### Competing Interest Statement

The authors have declared no competing interest.

