## Supplemental Tables for "Phylosymbiosis and the hologenome in fungus-gardening ants"

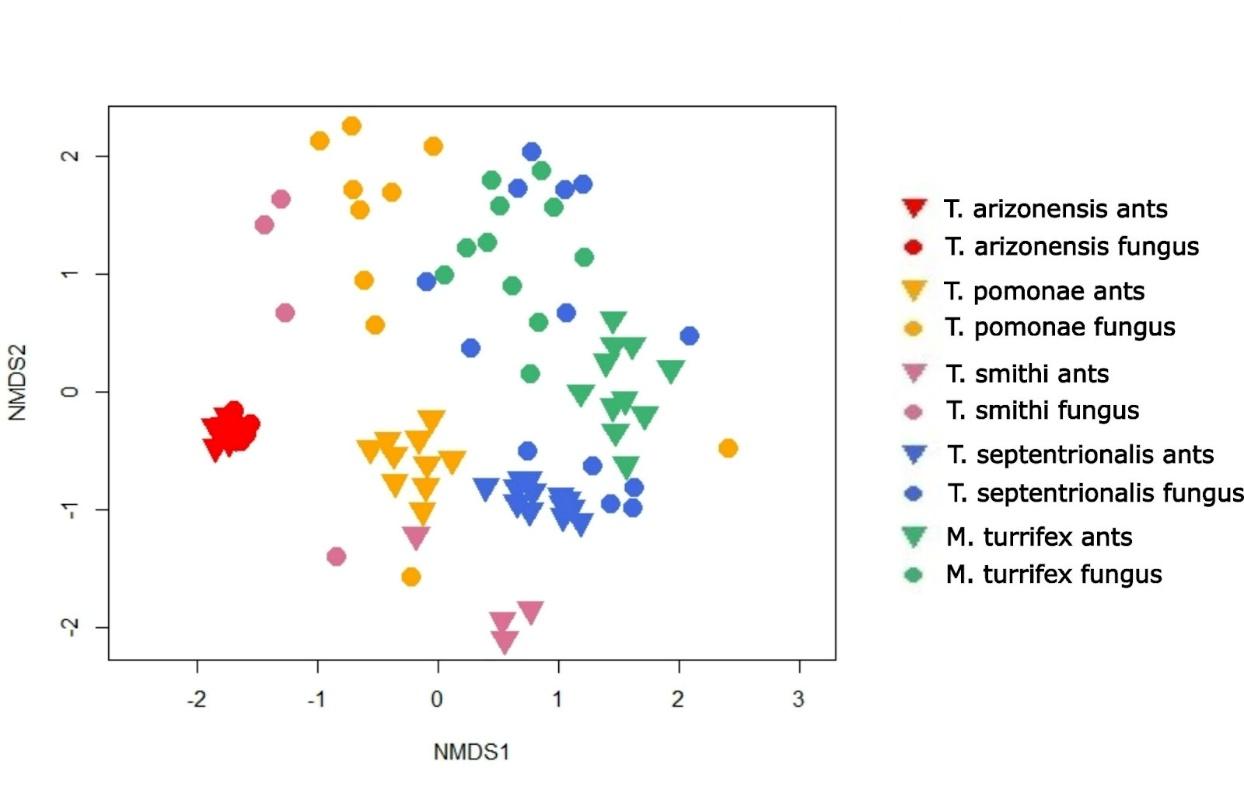

**Figure S1** Non-metric dimensional scaling (NMDS) plot depicting similarity between ant-associated microbiome and fungus-associated microbiome samples for *Mycetomoellerius turrifex, Trachymyrmex arizonensis*, *T. pomonae*, *T. septentrionalis*, and *T. smithi*. Ants and fungi of each species are clustering together, which indicates overall similarity. Species are represented by different colors whereas ants and fungi are represented by triangles and circles, respectively.

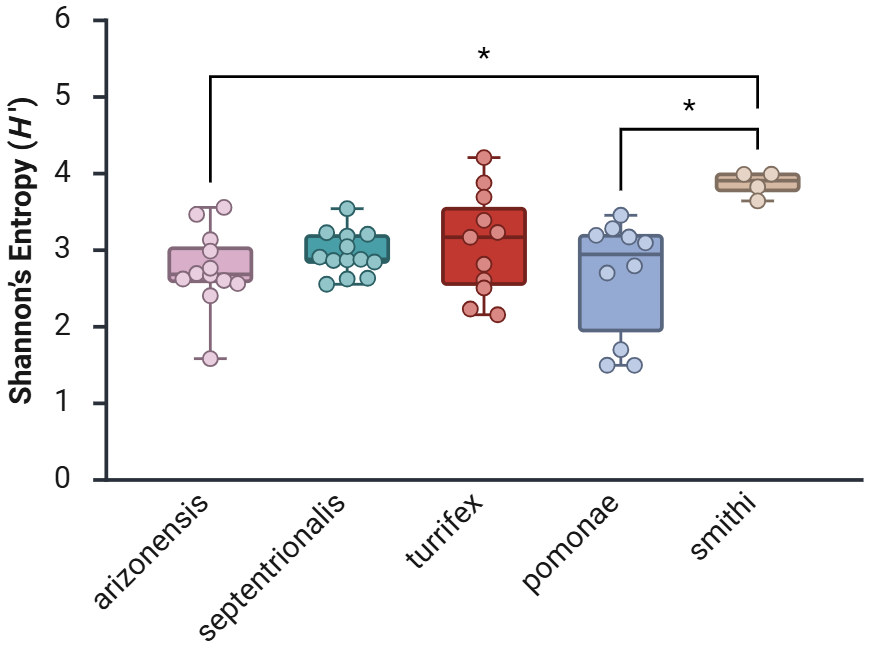

Figure S2. Shannon’s Entropy values of alpha diversity among all ant-associated microbiomes by ant species. Lines correspond to significant Dunn’s tests. *T. arizonensis* ant-associated bacterial microbiomes were significantly lower than the community found in *T. smithii*, which was significantly higher than *T. pomonae* (Kruskal Wallis Test, df =4, χ2=11.191, p-value=0.0245).

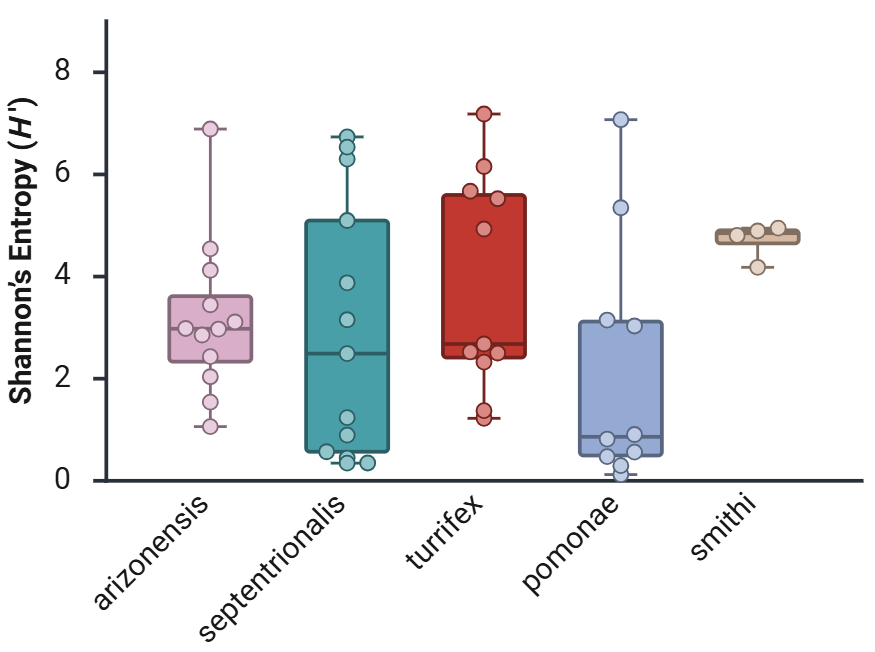

Figure S3. Shannon’s Entropy values of alpha diversity among all fungus-associated microbiomes by ant species. Bacterial diversity of fungus gardens did not significantly differ across all ant species. (Kruskal Wallis Test, df=4, χ2=6.091, p-value=0.1925

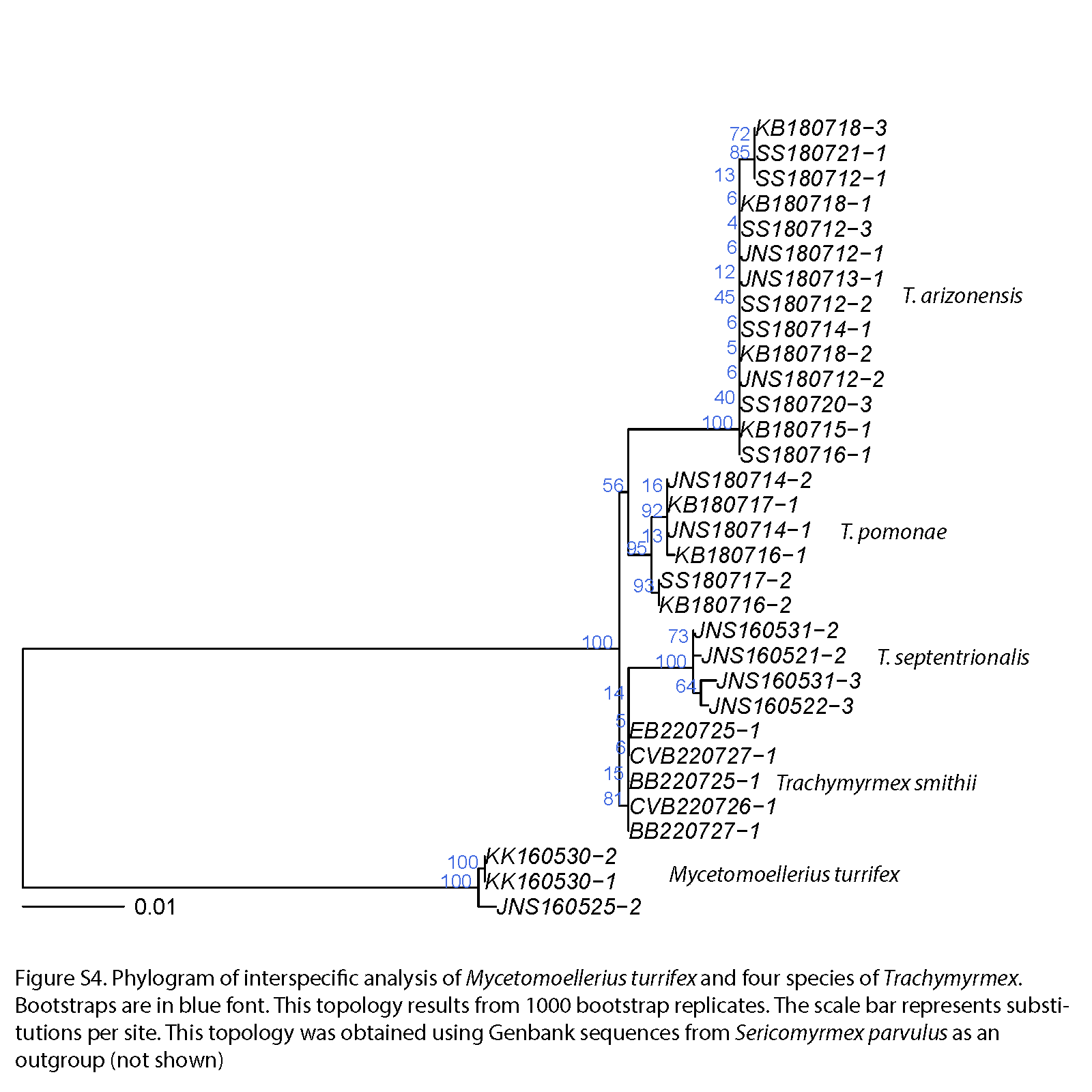

Figure S4. Phylogram of interspecic analysis of *Mycetomoellerius turrifex* and four species of *Trachymyrmex*. Bootstraps are in blue font. This topology results from 1000 bootstrap replicates. The scale bar represents substitutions per site. This topology was obtained using Genbank sequences from *Sericomyrmex parvulus* as an outgroup (not shown).

Table 1. See Excel spreadsheet that contains all collection locations and NCBI Accession numbers.

Table 2. P-values from Dunn’s tests on species comparisons of Ant and Fungal alpha diversity measures.

| **Ants** |  | |
| --- | --- | --- |
| Comparison | p-value | |
| *T. arizonensis vs T. septentrionalis* | 1 | |
| *T. arizonensis vs M. turrifex* | 1 | |
| *T. arizonensis vs T. pomonae* | 1 | |
| *T. arizonensis vs T. smithi* | 0.01336 | |
| *T. septentrionalis vs M. turrifex* | 1 | |
| *T. septentrionalis vs T. pomonae* | 1 | |
| *T. septentrionalis vs T. smithi* | 0.1093 | |
| *M. turrifex vs T. pomonae* | 1 | |
| *M. turrifex vs T. smithi* | 0.2274 | |
| *T. pomonae vs T. smithi* | 0.03815 | |
| **Fungi** | |  |
| Comparison | | p-value |
| *T. arizonensis vs T. septentrionalis* | | 1 |
| *T. arizonensis vs M. turrifex* | | 1 |
| *T. arizonensis vs T. pomonae* | | 1 |
| *T. arizonensis vs T. smithi* | | 1 |
| *T. septentrionalis vs M. turrifex* | | 1 |
| *T. septentrionalis vs T. pomonae* | | 1 |
| *T. septentrionalis vs T. smithi* | | 1 |
| *M. turrifex vs T. pomonae* | | 0.6601 |
| *M. turrifex vs T. smithi* | | 1 |
| *T. pomonae vs T. smithi* | | 0.3384 |
